## Supplemental Figures for "Beyond BioID: Streptavidin outcompetes antibody fluorescence signals in protein localization and readily visualises targets evading immunofluorescence detection"

### Figure S1

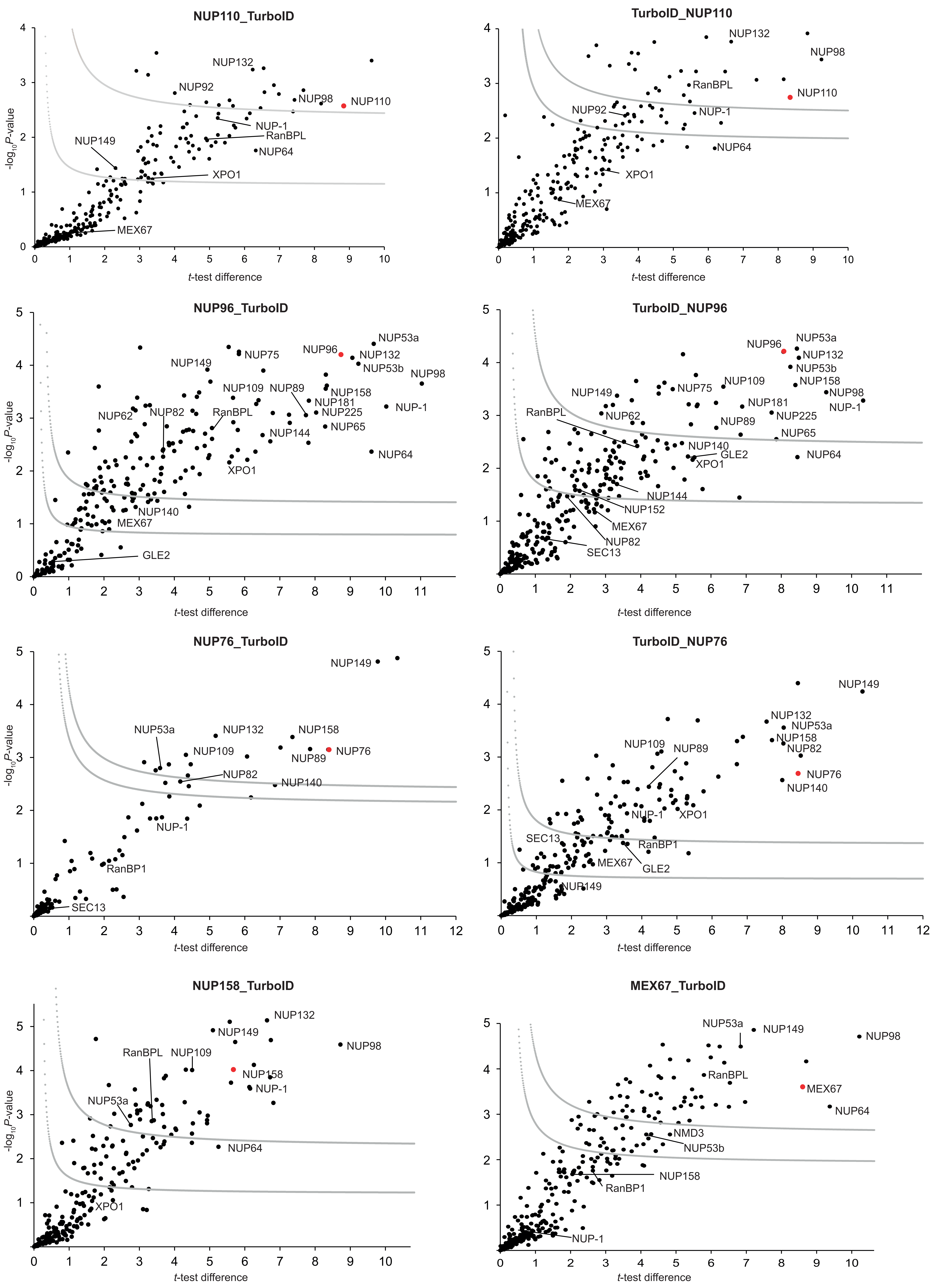


**Figure S1: Statistical analysis of NUP TurboID experiments.** Hawaii plot (multiple volcano plots) of LFQ results of the BioID experiments for NUP110, NUP96 and NUP76 with fused TurboID-HA tag either at the N- or C-terminus, and C-terminally tagged NUP158 and MEX67. All samples were prepared at least in duplicate. To generate the volcano plots, the −log10*P*-value was plotted versus the *t*-test difference (difference between means), comparing each respective bait experiment to the wt control. Potential interactors were classified according to their position in the plot, applying cutoff curves for "significant class A" (SigA; gray, upper curve; FDR = 0.01, s0 = 0.1) and "significant class B" (SigB; gray, lower curve; FDR = 0.05, s0 = 0.1), respectively. Bait proteins are indicated by a red dot, known NUPs and transport factors (as shown in Figure 1A) are labelled and LFQ data is given in Table S2.

Figure S2

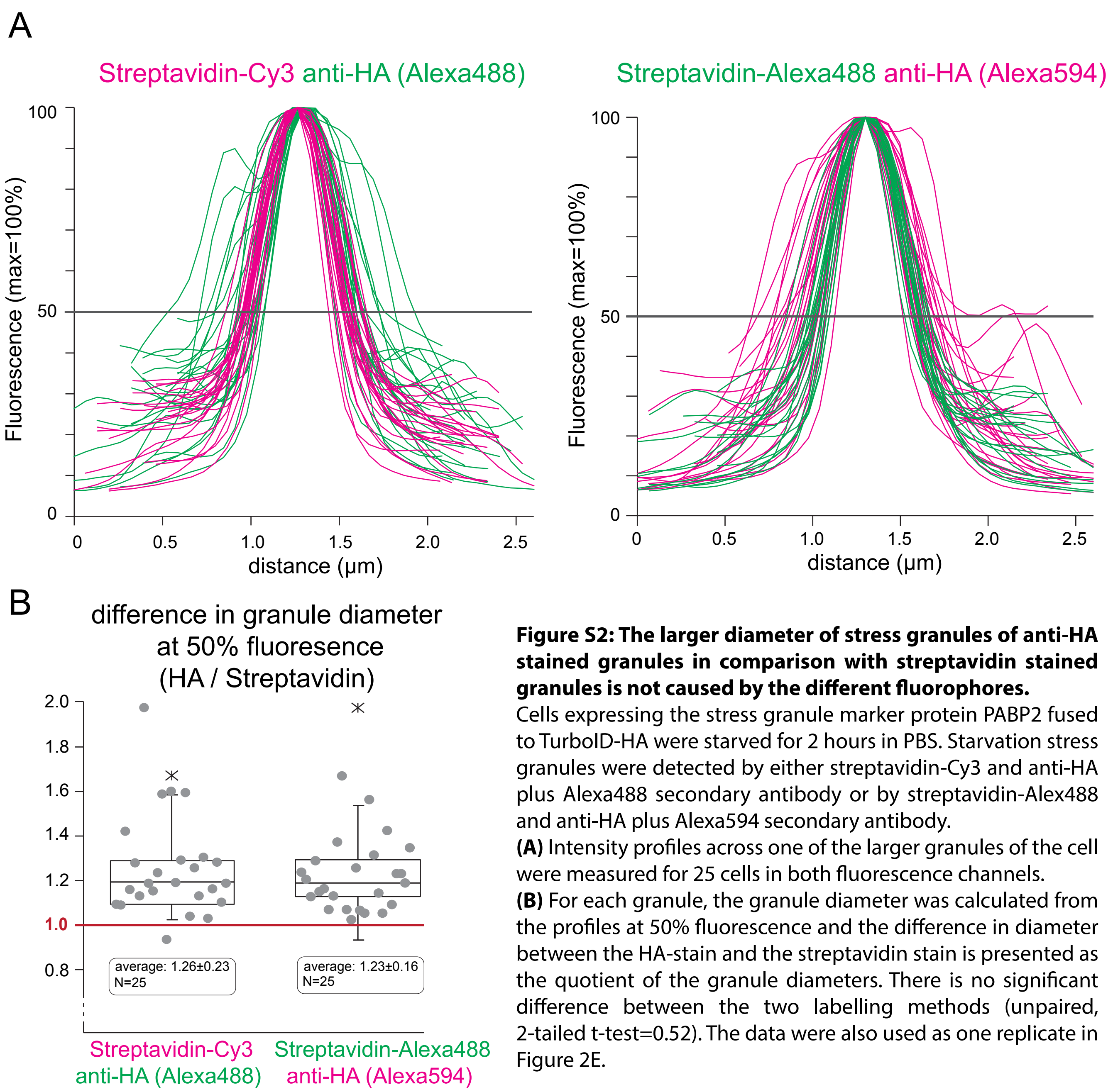


**Figure S2: The larger diameter of stress granules of anti-HA stained granules in comparison with streptavidin stained granules is not caused by the different fluorophores.**
Cells expressing the stress granule marker protein PABP2 fused to TurboID-HA were starved for 2 hours in PBS. Starvation stress granules were detected by either streptavidin-Cy3 and anti-HA plus Alexa488 secondary antibody or by streptavidin-Alex488 and anti-HA plus Alexa594 secondary antibody.
**(A)** Intensity profiles across one of the larger granules of the cell were measured for 25 cells in both fluorescence channels.
**(B)** For each granule, the granule diameter was calculated from the profiles at 50% fluorescence and the difference in diameter between the HA-stain and the streptavidin stain is presented as the quotient of the granule diameters. There is no significant difference between the two labelling methods (unpaired, 2-tailed t-test=0.52). The data were also used as one replicate in Figure 2E.

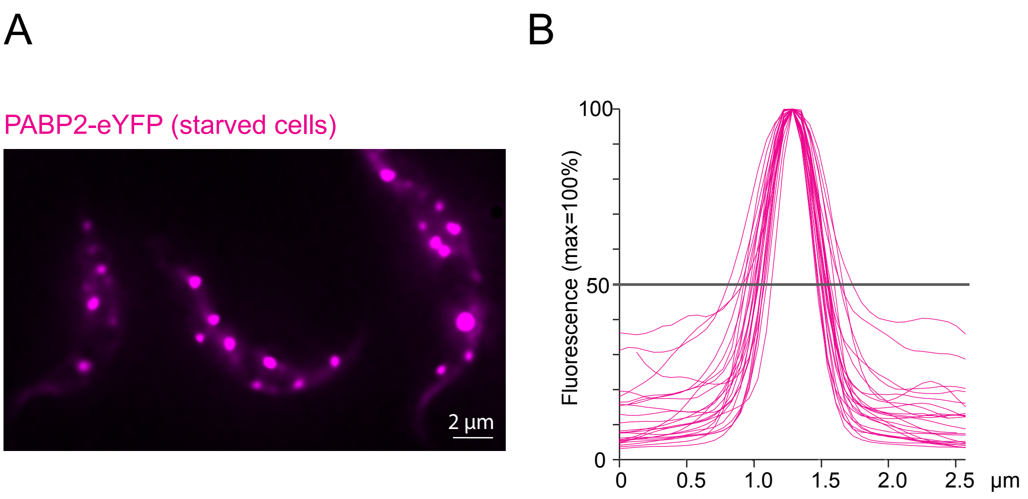


**Figure S3: PABP2-eYFP is not at the periphery of stress granules**
PABP2 was expressed as a C-terminal fusion to eYFP from the endogenous locus.
**A)** Fluorescence microscopy image of starved trypanosome cells (2 hours PBS) expressing PABP2-eYFP. The eYFP granules appear dot-like with a clearly defined shape, similar to the granules labelled with streptavidin in PABP2-TurboID-HA expressing cells.
**B)** The fluorescence intensity profile is shown for 25 granules. The profiles are similar to the profiles of the streptavidin labelled granules from PABP2-TurboID-HA cells but differ from the profiles obained with anti-HA (compare Figure 2). The shape of the profiles is consistent with an even distribution of PABP2 throughout the (spherical) stress granules, rather than a peripheral localisation.

Figure S4

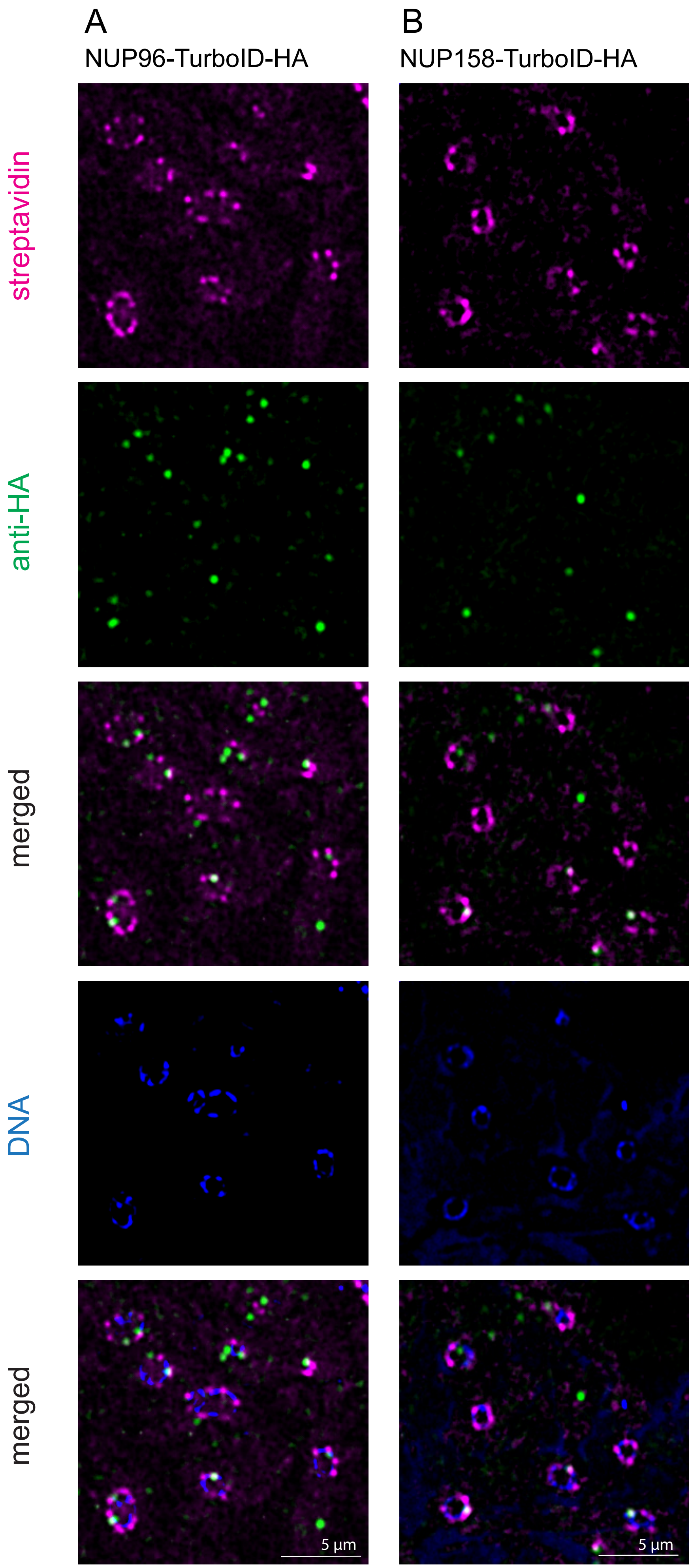


**Figure S4: Streptavidin and anti-HA signal on LR-white embedded sections**

Trypanosome cells expressing NUP96-TurboID HA (A) or NUP158-TurboID-HA (B) were high-pressure frozen and embedded in LR-white. The resin with the embedded cells was cut into 100 nm tick slices and the proteins were detected by streptavidin and anti-HA on the surface of these slices. Streptavidin labels significantly more nuclear pores than anti-HA.

Figure S5

A TurboID-HA-MLP2

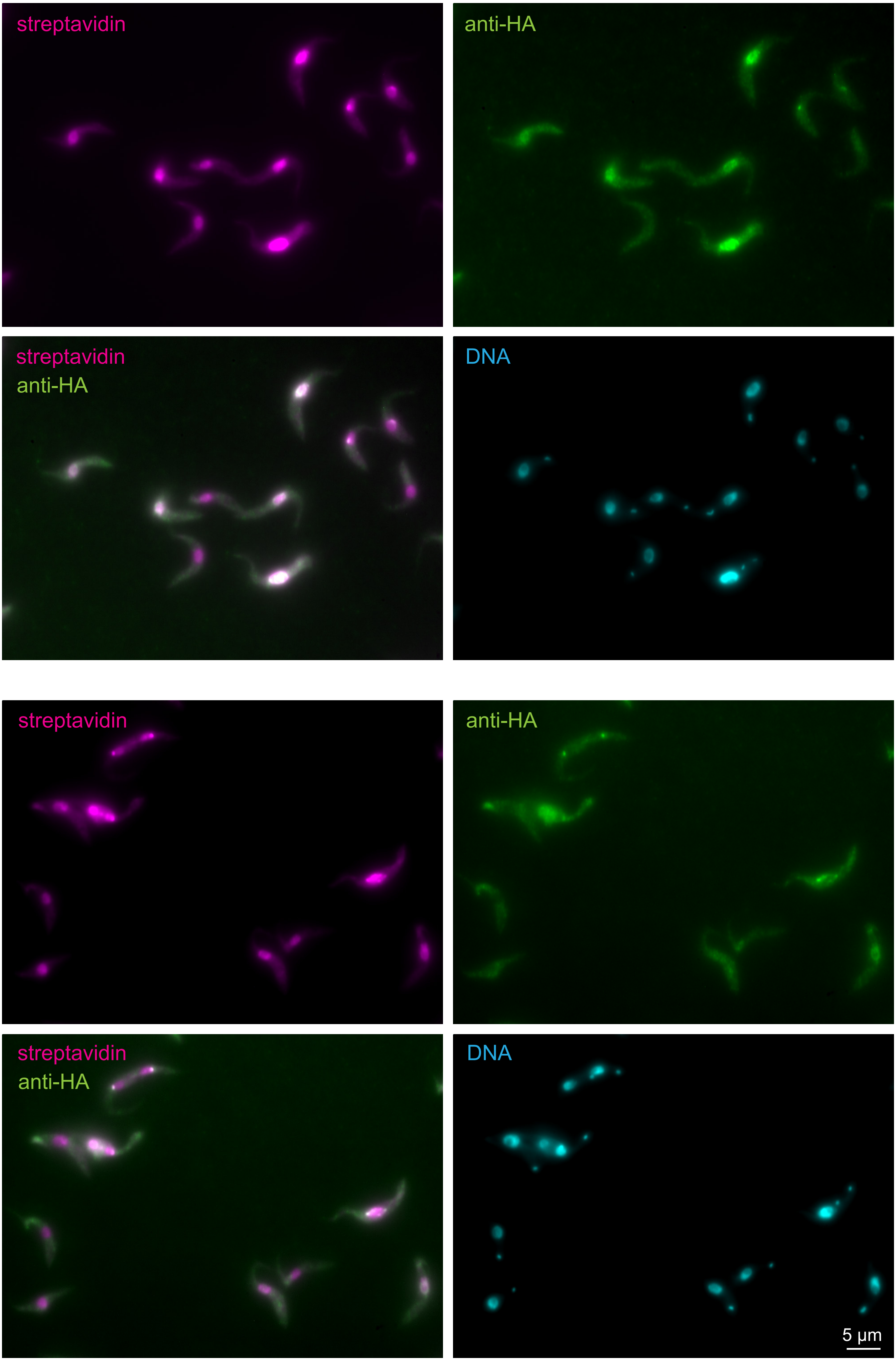

Figure S5

B MLP2-TurboID-HA

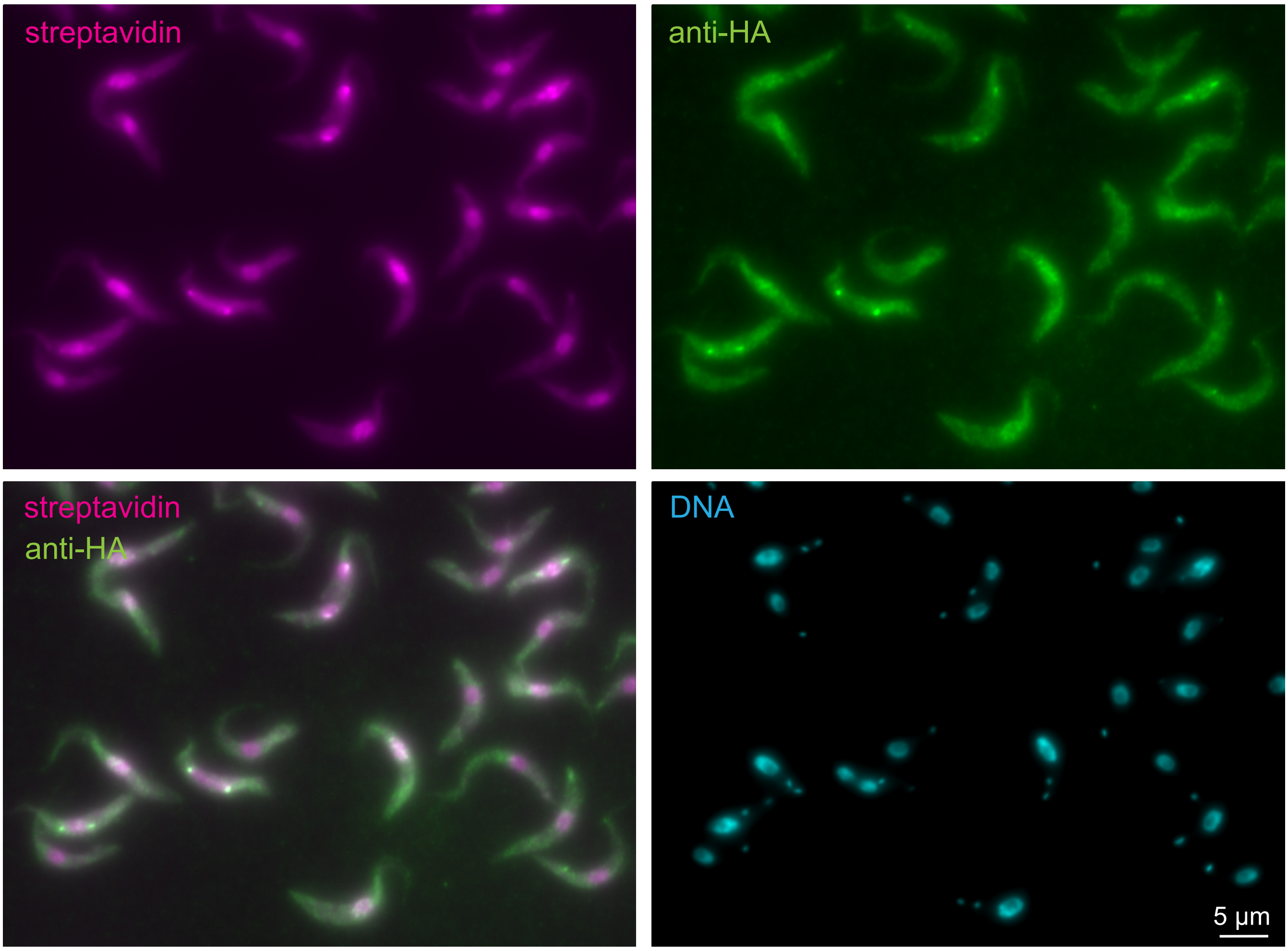


**Figure S5: MLP2 does largely not localise to nuclear pores, but to the nucleus and to the spindle pole.**
*T. brucei* MLP2 was expressed fused to either an N-terminal TurboID-HA tag (A) or a C-terminal TurboID-HA tag (B) from the endogenous locus. The cells were labelled with cy3-streptavidin (pink) and anti-HA (green). Unprocessed images are shown as Z-stack projections (sum slices, 28 stacks).

### Figure S6 (page 1)

A

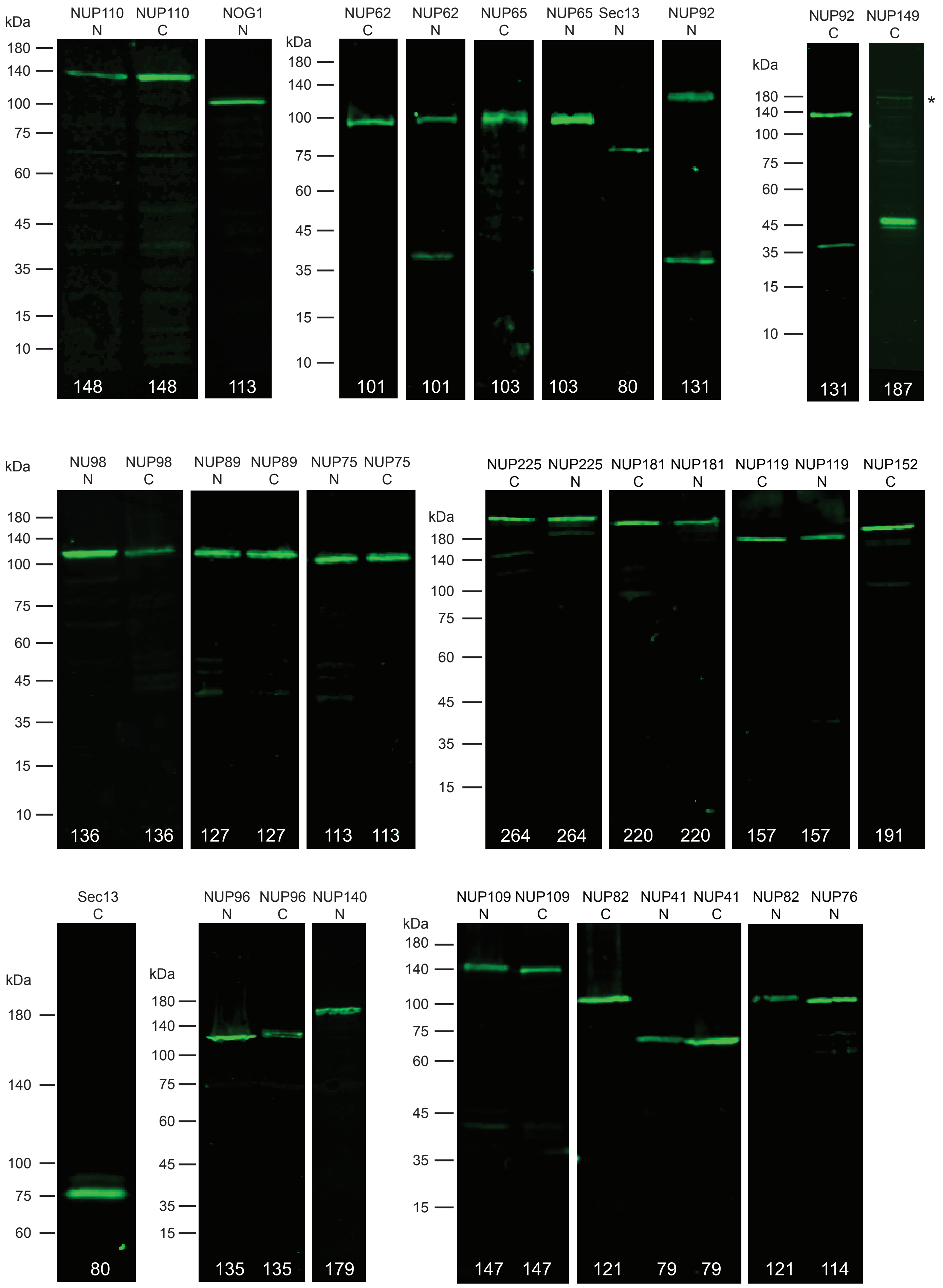

Figure S6 (page 2)

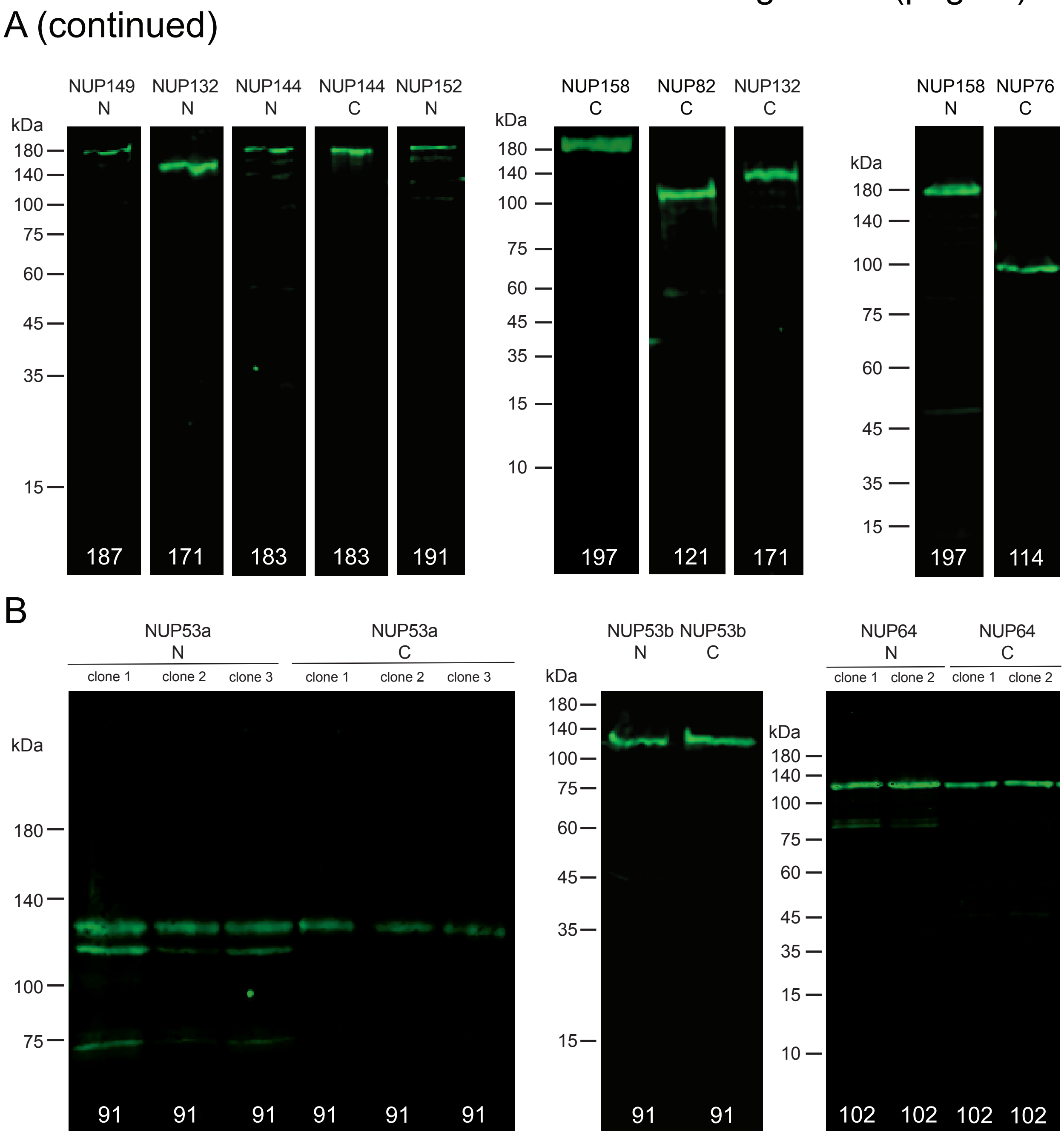


**Figure S6: Western blots**

Protein extracts of all cell lines expressing TurboID-HA fusion proteins (fused to either the N or C-terminus, as indicated) were loaded on SDS PAGE, gels blotted and the membranes probed with anti-HA. The expected molecular weight of the fusion protein is indicated in white numbers. **(A)** For most proteins, one single band at roughly the expected MW was detected; occassionally, there was a second smaller band, likely due to degradation. For C-terminally tagged NUP149 the smaller band was more prominent than the band at the correct MW (asteriks). **(B)** For the small FG-NUPs NUP53a, NUP53b and NUP64 we observed a larger MW than expected. However, this was repeatedly observed for independent transfections, for independent clonal cell lines and there was no difference between N- and C-terminally tagged version of the NUPs. We therefore assume that special features of these NUPs affect the migration behaviour on the SDS page.

### Supplementary Figure S7

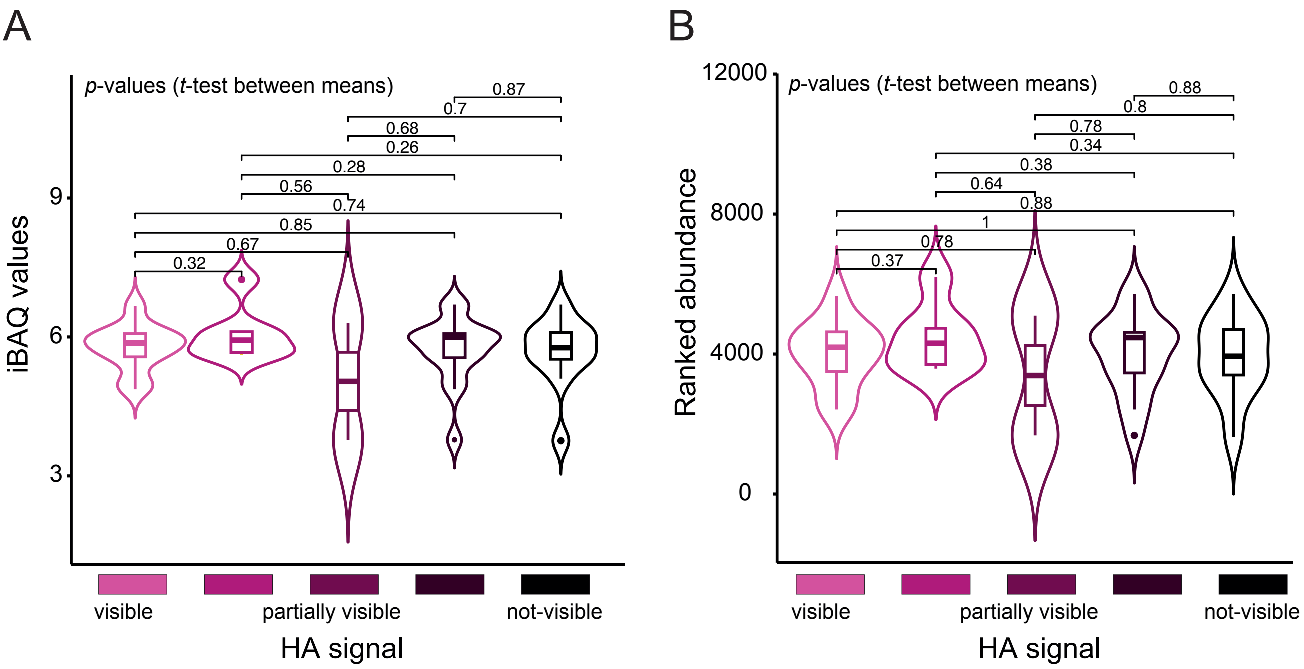


**Figure S7.** The strength of the immunofluorescence signal of HA-tagged nucleoporins does not correlate to estimated protein abundances. Estimation of protein abundances is available as intensity-based quantification (iBAQ) and ranked-order relative protein abundance values for the proteome of the *T. brucei* 927 strain [1]. Immunofluorescence signal of HA-tagged nucleoporins was categorized based on signal intensity as shown in Figure 5. We compared the means of protein abundances in iBAQ **(A)** and ranked-order relative abundance **(B)** between these categories using a t-test. A box plot was merged with a violin plot to show the distribution of the data points using the ggpubr package in R. The p-values of comparisons between all categories are indicated.

[1] M. Tinti and M. A. J. Ferguson, "Visualisation of proteome-wide ordered protein abundances in Trypanosoma brucei," Wellcome Open Res, vol. 7, p. 34, Feb. 2023, doi: 10.12688/wellcomeopenres.17607.2.

Figure S8

A *T. brucei* wild type cells

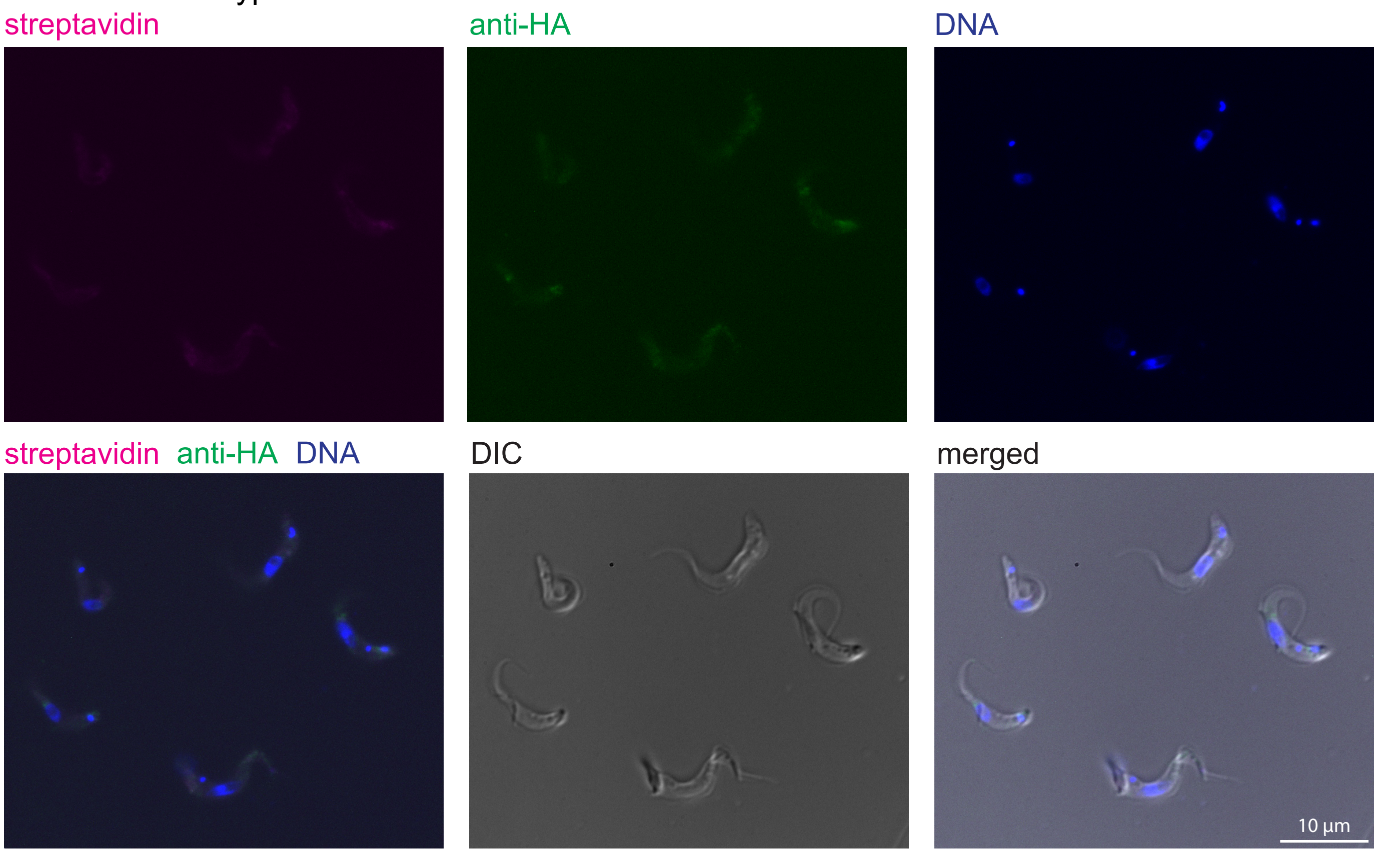


B HeLa cells, upper cell TurboID-HA-Nup88, lower cell wild type (not transfected)

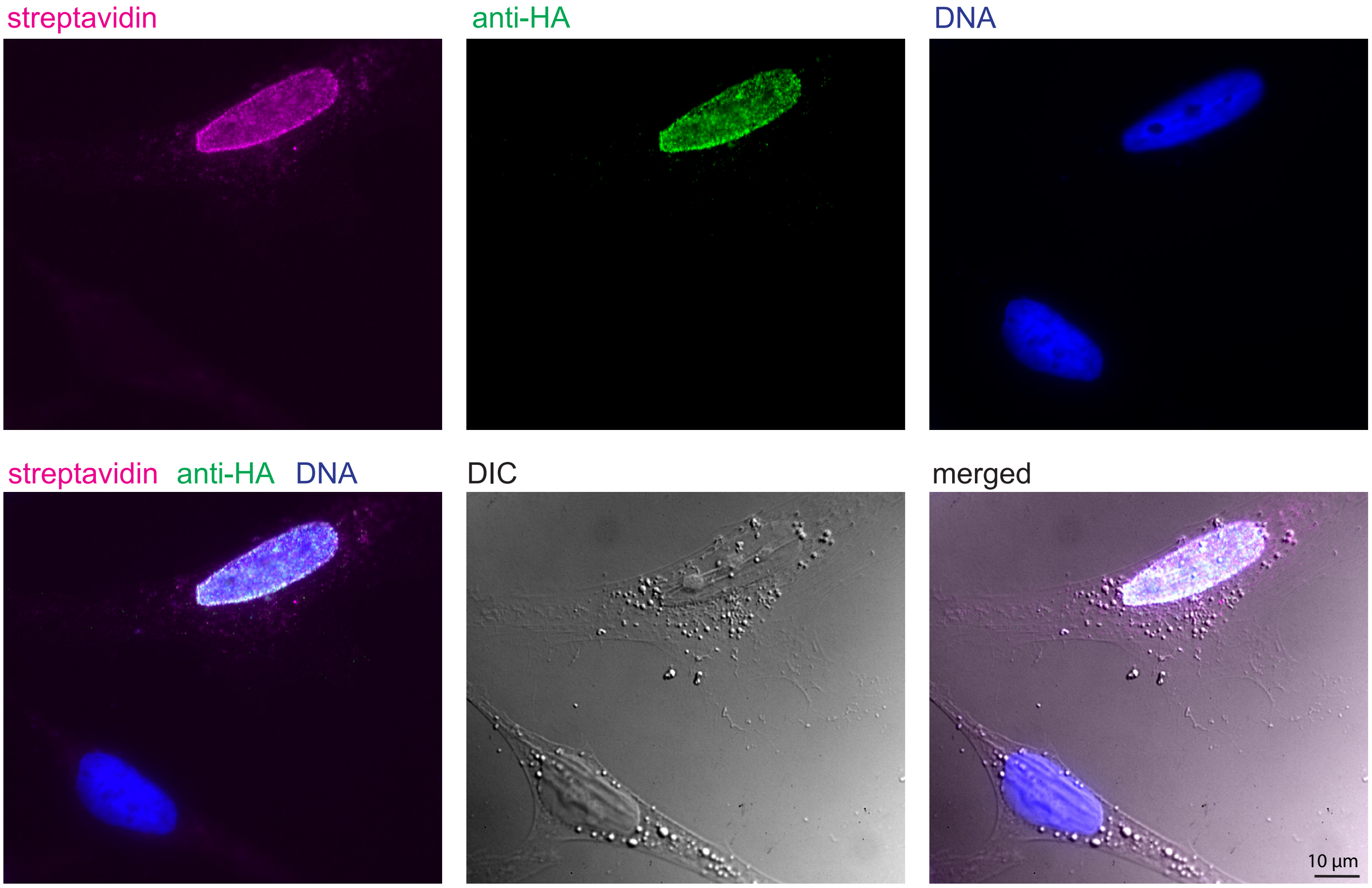


**Figure S8: Streptavidin and anti-HA signal of wild type cells**
**(A)** *T. brucei* wild type cells were labelled with streptavidin and anti-HA. A single plane image is shown.
**(B)** Streptavidin imaging of HeLa cells: the upper cell is transfected with TurboID-HA-NUP88, the lower one is not.
